# Cooperative Membrane Binding of HIV-1 Matrix Proteins

**DOI:** 10.1101/2023.09.22.559012

**Authors:** Puja Banerjee, Viviana Monje-Galvan, Gregory A. Voth

## Abstract

The HIV-1 assembly process begins with a newly synthesized Gag polyprotein being targeted to the inner leaflet of the plasma membrane of the infected cells to form immature viral particles. Gag-membrane interactions are mediated through the myristoylated(Myr) N-terminal matrix (MA) domain of Gag which eventually multimerize on the membrane to form trimers and higher-order oligomers. The study of the structure and dynamics of peripheral membrane proteins like MA has been challenging for both experimental and computational studies due to the complex dynamics of protein-membrane interactions. Although the roles of anionic phospholipids (PIP2, PS) and the Myr group in the membrane targeting and stable membrane binding of MA are now well-established, the cooperative interactions between MA monomers and MA-membrane still remain elusive. Our present study focuses on the membrane binding dynamics of a higher-order oligomeric structure of MA protein (a dimer of trimers), which has not been explored before. Employing time-lagged independent component analysis (tICA) to our microsecond-long trajectories, we investigate conformational changes of the matrix protein induced by membrane binding. Interestingly, the Myr switch of a MA monomer correlates with the conformational switch of adjacent monomers in the same trimer. Together, our findings suggest that MA trimerization facilitates Myr insertion, but MA trimer-trimer interactions in the lattice of immature HIV-1 particles can hinder the same. Additionally, local lipid density patterns of different lipid species provide a signature of the initial stage of lipid-domain formation upon membrane binding of the protein complex.

**TOC:** 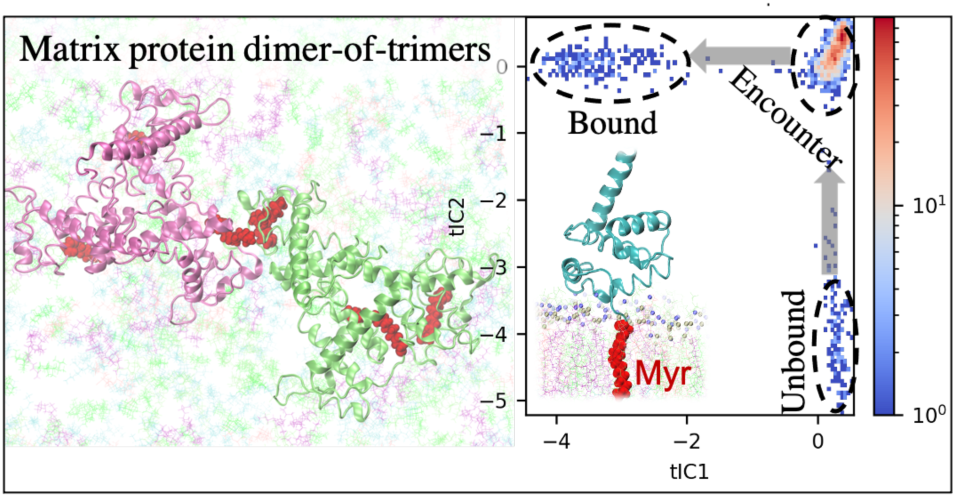

## Introduction

The human immunodeficiency virus type 1 (HIV-1) assembly process is mediated primarily by the retroviral group-specific antigen (Gag) polyprotein ^1, 2^. Membrane binding of Gag to the inner leaflet of the plasma membrane (PM) of the host cell is a crucial step in the assembly process. The Gag polyprotein consists of the matrix (MA), Capsid (CA), spacer-peptide 1 (SP1), nucleocapsid (NC), spacer-peptide 2 (SP2), and p6 domains. While the CA and NC domains contribute to Gag multimerization and RNA encapsidation during the viral assembly ^3–5^, the p6 domain plays an important role in viral budding. Membrane targeting to the PM and recruitment of the Envelope(Env) protein is mediated by the MA domain, which is post-translationally acylated with a myristate (Myr) group in its N-terminus ^6–10^. Structural studies have shown that MA proteins primarily oligomerize into trimeric structures, which then undergo higher-order oligomerization to form a lattice of hexamer-of-trimers ^11^. MA binding to the PM is facilitated by several key factors, including electrostatic interactions of its highly basic region (HBR) with negatively charged lipid headgroups, hydrophobic interactions of the Myr group, membrane internal structure, etc. ^11–20^. Moreover, myristoylated MA ((+)Myr-MA) prefers binding specifically to membrane microdomains enriched in PI(4,5)P_2_ and cholesterol ^21, 22^. The relative contribution of these MA-lipid interactions is nontrivial to determine; however, it likely depends on the multimerization state of Gag and the specific protein-protein interface.

MA protein is a peripheral membrane protein (PMP) whose biological activity is predominantly determined by its ability to switch between a soluble and membrane-bound state. Unlike integral membrane proteins, specific protein-lipid binding site structures of PMPs are hard to determine both for experiment and simulation studies due to the unique nature of PMP-membrane interactions, which are often transient ^12, 18, 23–29^. A few long-timescale all-atom molecular dynamics (AAMD) simulations aimed to explore this non-equilibrium process of PMP binding with membrane ^30–35^. Although the strategy of AAMD simulations to study the membrane binding of PMP seems simple, it is computationally expensive even for a small monomeric PMP. Moreover, the HIV-1 MA assembly process is more complex as MA binding and MA aggregation also modify the structure and dynamics of the membrane.

The PM of eukaryotic cells is asymmetric in lipid distribution and lipid unsaturation. The inner PM leaflet is enriched in unsaturated phospholipids, like phosphatidylethanolamine (PE), phosphatidylserine (PS), phosphatidylcholine (PC), a small amount of phosphatidylinositol (PI) lipids, and cholesterol. Whereas the outer PM leaflet mainly contains phosphatidylcholine (PC), saturated sphingomyelin lipids (SM), and cholesterol. Cholesterol helps counteract unfavorable packing of saturated and unsaturated lipids in membranes in general and maintains structural integrity and membrane fluidity. Experimental studies on HIV-1 assembly dynamics have suggested that Gag multimerization on the PM induces clustering of specific lipid species like PIP2 and cholesterol in the inner leaflet, which in turn remodels the outer leaflet through trans bilayer coupling ^15, 36–39^. The relationship between MA multimerization and membrane binding is still unclear as well as its impact on the lateral organization of lipids in each PM leaflet.

A cryo-electron tomography (cryo-ET) experimental study by Briggs and coworkers has resolved the MA trimer-trimer interacting interface in immature virus particles ^12^. Based on the cryo-ET density map, they suggested no Myr group remains in the sequestered position in the immature MA lattice (**Figure 1**). MA trimer-trimer interactions in the fitted atomic model (PDB ID: 7OVQ) are mediated by N-terminal residues, namely α-helix 1(H1) and 3_10_ helix (shown in **Figure 1E**). Another experimental study by Saad and coworkers studied such MA-MA interactions in crystal structures using X-Ray crystallography and NMR spectroscopy ^20^. The effect of this kind of MA-MA interaction on membrane binding and membrane reorganization is the focus of the present study.

**Figure 1:**
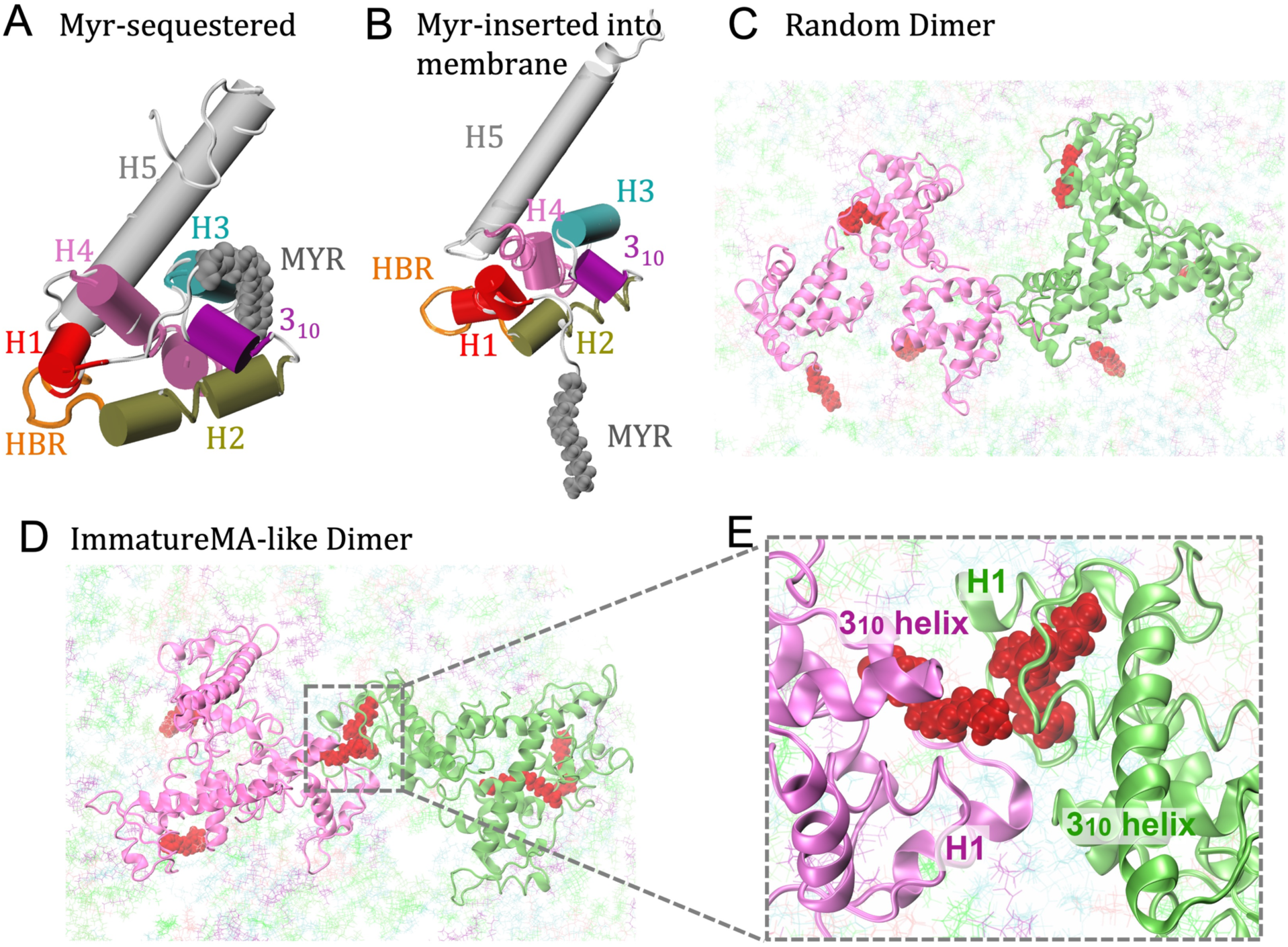
Side-view of the membrane-bound MA monomers in two different states of Myr switch. Key regions are labeled: H1, HBR, H2, 3_10_ helix, H3, H4, and H5 (in order of appearance in the protein sequence) (A) Myr group is sequestered into the hydrophobic pocket and (B) Myr group is inserted into the membrane. (C-D) Top-view of the final structures of MA dimer of trimers at 5 μs of simulation. (E) PPI in the trimer-trimer interface of immature MA-like dimer.

In a previous AAMD study, we characterized the membrane binding mechanism of MA monomers and pre-formed trimers, along with details of Myr insertion and the initial membrane response to protein binding ^31^. It was observed that the trimer conformation enhances lipid rearrangement and accelerates Myr insertion. An enrichment of PIP2 at the MA-binding site has been suggested several times before ^11, 40, 41^. Although it is well-accepted now that MA trimerization and PIP2 recruitment enhance the Myr exposure ^10, 18^, the effect of MA trimer-trimer interactions on Myr insertion dynamics in a membrane-bound MA protein complex is yet to be explored.

We performed and reported here microsecond-long AAMD simulations with two MA trimers initially placed above an asymmetric membrane model. Our realistic membrane model is based on lipidomic analysis of HIV-1 particles produced in HeLa cells reported by Lorizate *et al* ^36^. Using different initial configurations of two formed MA trimers, we examine the effect of MA-MA interactions on protein binding dynamics. Myr insertion dynamics is observed to be correlated with MA-MA interactions. Finally, the simulated data reveals enhancement of domain-forming lipids in the outer leaflet at regions that correspond to the protein binding site on the inner leaflet.

## Results

### Membrane binding and Myr insertion of matrix (MA) trimer assembly

We have carried out AAMD simulations with a pair of myristoylated-MA (Myr-MA) trimers in proximity to an asymmetric bilayer. To mimic the plasma membrane environment during HIV-1 assembly, we have carefully chosen the lipid composition of our membrane model. The inner leaflet is enriched with PS, PIP2, and PE lipids while the outer leaflet contains SM and cholesterol (for details, see ***Materials and Methods*** section). Depending on the initial arrangement of two MA trimers we have simulated two different systems, each for 5 μs, to investigate Myr insertion of the MA trimers and the role of MA trimer-trimer interactions on that. Two systems are shown in ***Figure 1***: (a) *Immature MA-like dimer*: MA trimeric interface is stabilized by the interactions of N terminal residues, α-helix 1 (H1) and 3_10_ helix, as seen in the immature virus particles ^12^ (***Figure 1E***). Initially, the MA dimer of trimers is kept separated from the asymmetric membrane by at least 1 nm. (b) *Random dimer*: At t = 0, two MA trimers are kept separated from each other at a random arrangement near the asymmetric membrane. We have analyzed detailed trimer-trimer interactions in these systems, which will be discussed later. The structures of membrane-bound MA monomers before and after the Myr switch are shown in ***Figure 1A-B*** highlighting key regions (5 αhelices, HBR domain, and 3_10_ helix). Domains are rearranged among themselves and αhelix structures are perturbed depending on the Myr group position, whether it is sequestered into the hydrophobic pocket of MA or inserted into the membrane. Furthermore, with our unbiased AAMD simulations, we could study spontaneous MA conformational change induced by membrane binding.

A previous AAMD study by Monje-Galvan *et al.* reported membrane insertion events of the Myr moiety of HIV-1 matrix protein in detail ^31^. According to that study, for a single MA monomer, Myr insertion takes place only from the ‘*open bound conformation*’ (helix1 and HBR domain of the MA interact with the membrane) when the domain between the Lid (3_10_ helix) and H1 opens up. The first carbon of Myr tail, C2 binds the membrane first and the rest of the tail explores the membrane surface for accessible large-enough lipid packing defect suitable for insertion. It was concluded that Myr insertion never occurs in a diving fashion, where the last carbon of the Myr tail, C14 inserts first ^31^. Although this mechanism of Myr insertion appears to be valid in our current simulations, the conformational exploration step is expected to be influenced by MA-MA interactions present in the system.

Our simulated data in this work suggest that MA trimer-trimer interactions (as observed in immature virions) can inhibit Myr insertion. ***Figure 2*** shows membrane binding events in the two different systems of interest. Within 5 μs, only 2 Myr groups insert into the membrane in the immature MA-like dimer model. None of the Myr groups at the trimer-trimer interface exhibit Myr insertion although they are in ‘*Myr-exposed*’ states. On the other hand, in the random dimer model (***Figure 2***), 4 Myr groups are at a stable inserted state at the end of the 5 μs trajectory. In both simulations, at least one Myr group remained in the sequestered state. The Myr insertion pattern in multiple independent trajectories of these systems suggests that the Myr switch is more feasible in the random trimer model for two probable reasons: (1) MA trimers are able to move independently on the membrane surface searching for lipid defects for the Myr insertion, (2) NTD conformations of MA proteins are not restrained due to MA trimer-trimer interactions. Thus, our findings reveal a competition between MA trimer-trimer interactions and the Myr switch of MA proteins.

**Figure 2:**
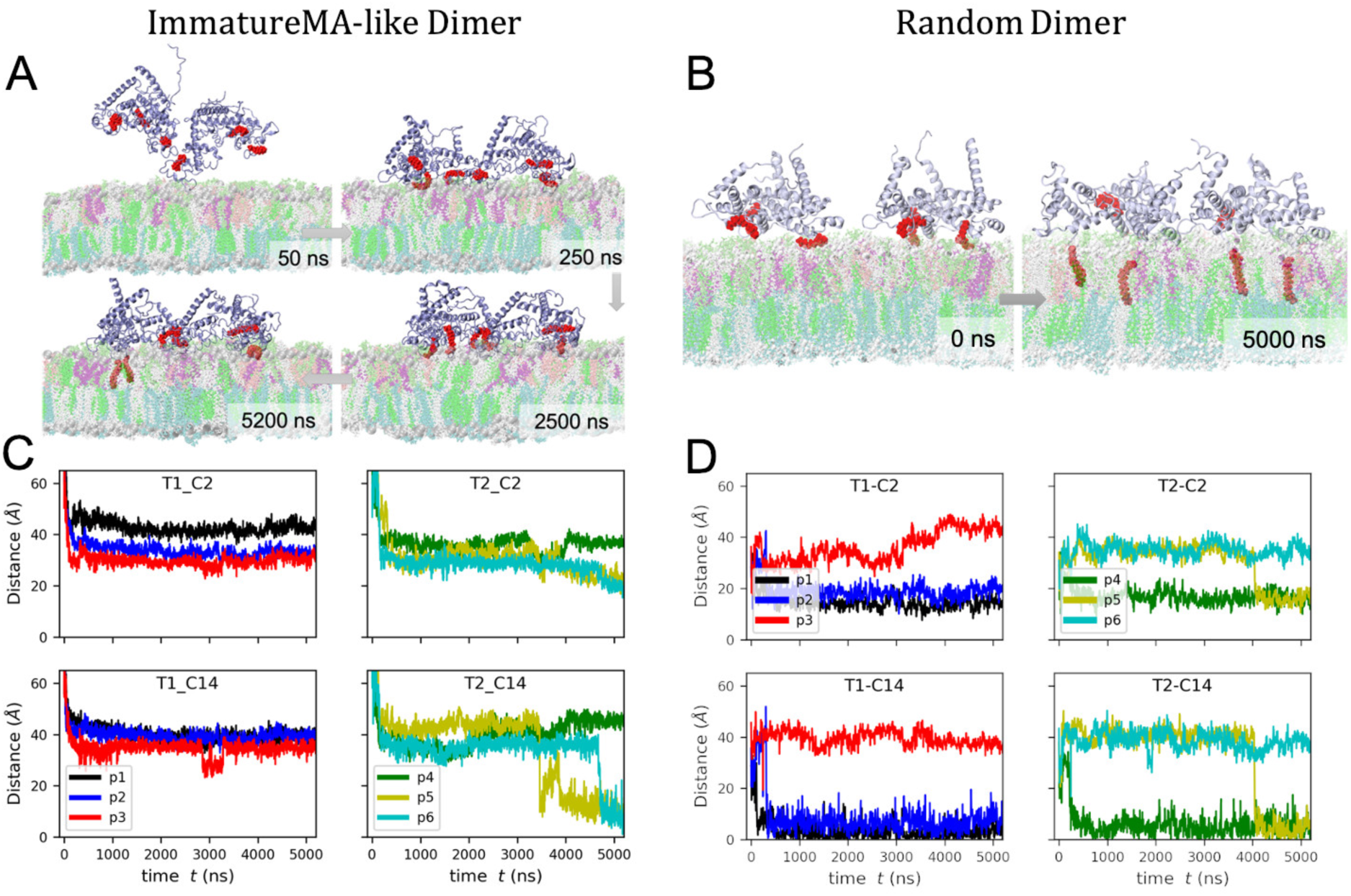
(A-B) Snapshots showing the progression of events for the immature MA-like and random dimer systems. (C-D) Time-evolution of the distance between the first carbon (C2) and the last carbon (C14) of the Myr group and the membrane center. Each protein unit is denoted as p1-p6, and each trimer as T1 and T2; p1, p2, p3 constitute Trimer1 and p4, p5, p6 belong to Trimer2.

MA trimer-trimer interface interactions in the two systems are quite different at the final step of the 5 μs trajectory (**Figure 3**). In the immature MA-like dimer, the MA trimer-trimer interface is stabilized by the interactions of N-terminal residues and the N terminus of helix 1 with themselves and the 3_10_ helices. The simulation started with this interface structure (where there were no MA-membrane contacts in the initial structure) and this MA-MA interface remained stable during membrane binding of the complex throughout the simulation. Therefore, this MA trimer-trimer interaction is observed to be not solely mediated by the membrane but stable in the solution phase also. However, in this scenario, protein-protein interactions (PPI) through N-terminal residues in the immature MA-like dimer hinder the conformational flexibility of the N-terminal domain (NTD) required for Myr insertion, as mentioned before.

**Figure 3:**
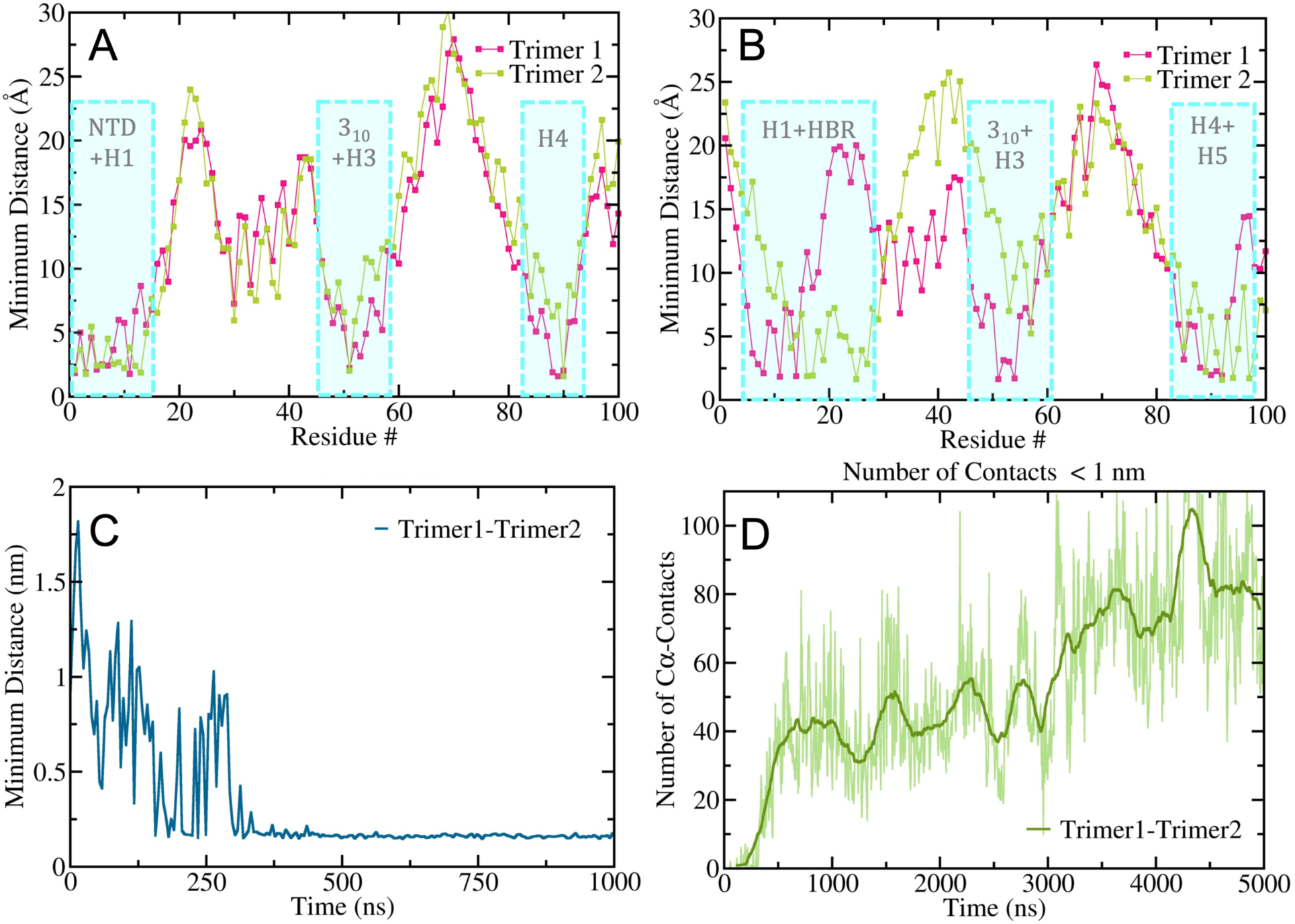
Identification of TTI in two MA complexes: Minimum distance of the adjacent trimer from the residues of trimer1 and trimer2 in (A) Immature MA-like dimer complex, (B) Random dimer complex. The final structures of 5 μs simulations are considered here. Binding domains are characterized in blue boxes. Time evolution of TTI during the assembly formation in the Random dimer model is monitored using the parameters: (C) minimum distance between two trimers, (D) number of Cα contacts (r_cut_=1 nm).

### MA trimer-trimer interactions (TTI) in two assembly structures

As mentioned earlier, we have carried out simulations with and without a stable trimer-trimer interface at the initial protein structures. In the random dimer model, starting from two separated membrane-bound MA trimers, trimer-trimer assembly formation occurs spontaneously in our simulations, driven by membrane and water-mediated interactions. However, in the Gag assembly process at the producer cell plasma membrane, CA SP1 assembly, IP6, and RNA binding promote and accelerate the MA assembly process. On the other hand, immature MA-like dimers maintained the interface during the membrane targeting and binding process in our simulated trajectories. This observation suggests that the dimerization of MA trimers in immature MA-like assembly is not purely lipid-mediated, which agrees with recent NMR studies by Saad and coworkers ^20^.

Here we have characterized TTI in two MA complexes. **Figure 3A-B** clearly distinguish between TTI for two different dimers of trimers, as seen at the final step of the 5 μs trajectory. In these figures, the minimum distance of residues of trimer1 and trimer2 from the other trimer are plotted against the residue numbers of the former. While, in the immature MA-like dimer model, (among others) N-terminal residues are involved in TTI (**Figure 3A**), in the random dimer model (among others) HBR domain residues are involved in TTI (**Figure 3B**). Also, unlike the immature MA-like dimer, the trimer-trimer interface in the random dimer system is not symmetric; two TTI profiles for trimer1 and trimer2 in **Figure 3B** are different. In the random dimer system, the progression of the MA trimer association process is investigated using the minimum distance between two trimers (**Figure 3C**) and the number of Cα contacts between trimer1 and trimer2 (for a cut-off radius of 1nm) (**Figure 3D**). While after ~ 300 ns, the assembled state is stable for random dimer complex, the trimer-trimer assembly samples over the different PPI networks and the number of contacts between two trimers keeps increasing throughout the trajectory.

### Membrane-induced conformational switch of MA monomers

Protein-protein interactions (PPI) within or on cellular membranes are often a key event in cellular signaling ^42, 43^. Prediction of PPI remains a challenging task for computer simulations as it is influenced by protein environments, such as macromolecular crowding, lipid molecules, and even the solvent environment ^44–47^, specific ion and buffer ^48, 49^. Therefore, it is expected that understanding PPI in a peripheral membrane protein complex is prohibitively difficult both from experiment and computational perspective. We first aimed to understand the conformational transition in each individual protein unit in the complex, and then we explored how membrane binding-induced structural change of protein impacts PPI in the MA multimeric complex or vice versa.

The membrane binding process results in protein conformational heterogeneity in different monomers of the MA complex depending on the Myr group position, either in the fully inserted configuration or sequestered in the hydrophobic pocket of the protein or at the membrane surface. Neutron reflectometry study confirms that the MA domain adopts different configurations on the membrane surface to aid the Myr group choose the proper orientation for the insertion ^29^. Six protein units with different conformations at the end of our 5 μs trajectory of immature MA-like dimer are shown in **Figure 4**. During membrane binding, prior to Myr insertion, helix1(H1) and HBR interact with the anionic membrane surface. However, in the initial binding phase of the MA complex, depending on the PIP2/PS lipid accessibility, protein-membrane interactions exhibit significant diversity. In p4-p6, HBR interacts closely with the membrane, whereas for p1-3, the MA-membrane interface looks different. The H2 helix plays an important role in membrane binding for these. Moreover, the conformations of H1 and H2 helices are perturbed in some of these protein units.

**Figure 4:**
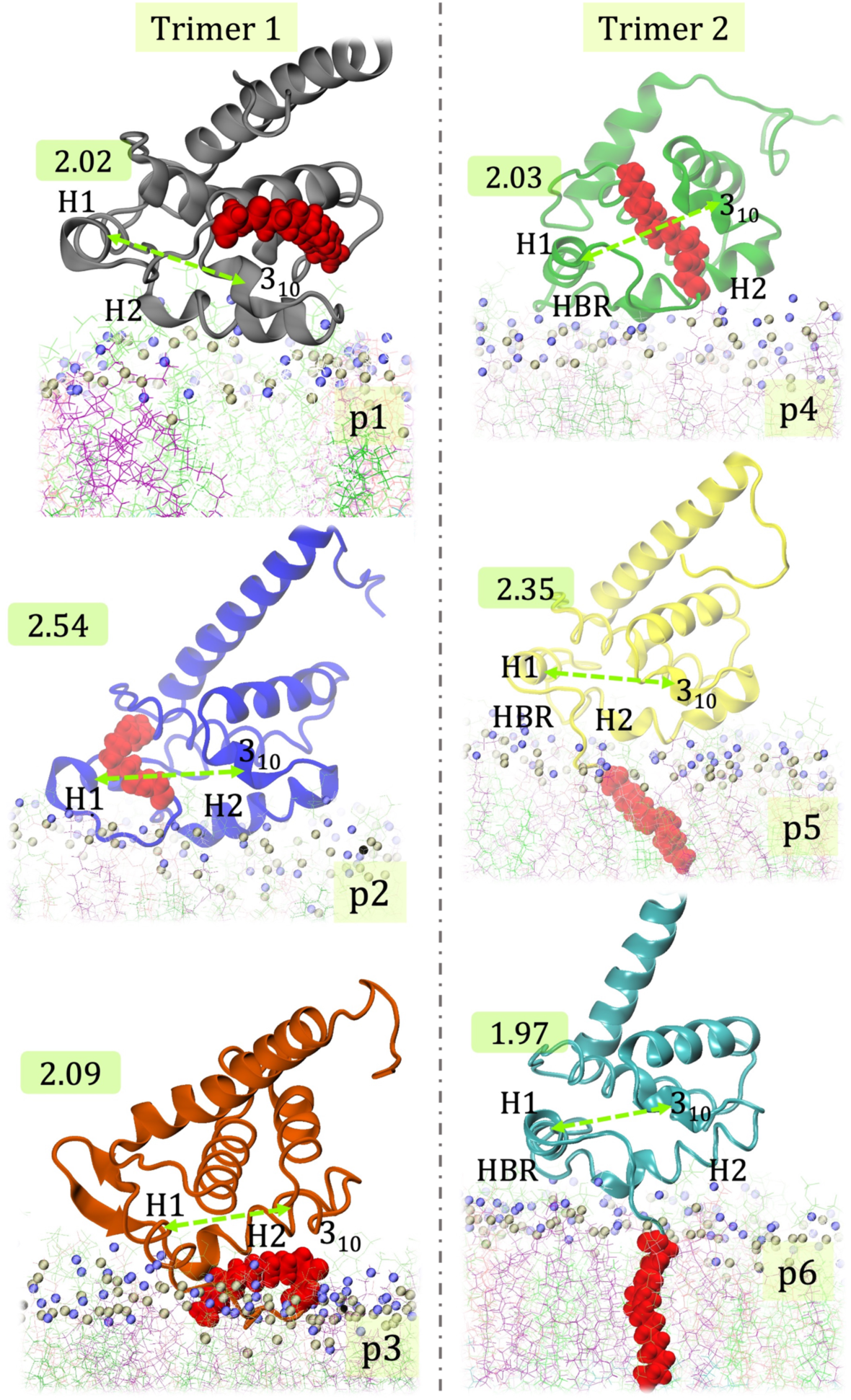
Structural heterogeneity of the six MA monomers (p1-6) at the end of the simulation trajectory of immature MA-like dimer. Distances between Helix1 and 3_10_ helices are highlighted by green dashed lines and the values (in nm) are shown in green boxes.

Fluctuation in the distance between the H1 helix and 3_10_ helix (d_H1-310_) plays an important role during Myr exposure ^31^. In the final structure of our simulation, d_H1-310_ for six monomers are observed to be ranging from 1.97 nm to 2.54 nm (Values of d_H1-310_(in nm) are shown in **Figure 4** in green boxes). For p6, in its stable Myr inserted state d_H1-310_ is minimum, whereas for p2, at an intermediate state of Myr insertion, this distance is maximum. Furthermore, the conformational transition in one protein unit is correlated with the dynamics of adjacent monomers. This aspect will be investigated later in more detail.

In our quest to understand the membrane binding mechanism of the MA-protein complex, we further looked into the structural transition of the protein units along the trajectory. Root mean-squared deviation (RMSD) and distance RMSD (DRMSD) analyses yield a direct measure of conformational fluctuation. Both are calculated with respect to the initial MA structure (at t = 0). Fluctuating domains like helix5 (H5) and N-terminal Myr are not considered in these calculations. DRMSD is calculated for the pair of atoms within 0.8 nm. In the immature MA-like dimer, protein unit 2(p2) of trimer1 and p4 of trimer 2 are at the trimer-trimer interface and the initial membrane anchoring of the protein complex is mediated by p2 around 30ns (**Figure 5**). RMSD analysis shows that p2 and the adjacent p3 monomer (RMSD > 0.4 nm) undergoes a structural transition at the beginning of the binding process, whereas most other monomers exhibit lower RMSD value throughout the simulation. As mentioned before, the p2 monomer first encounters the membrane and makes contacts with the anionic lipid molecules (PIP2, shown in green color) through its HBR domain. Although RMSD analysis could not differentiate the conformational change of the p2 monomer from the adjacent p3 monomer, pairwise DRMSD could capture the structural transition of the p2 monomer at the moment of its membrane anchoring (**Figure 5B**). The distance between the H1 helix and the 3_10_ helix increases for p2 at that point as it attains the ‘open conformation’ (**Figure 5D**), described in Ref. ^31^.

**Figure 5:**
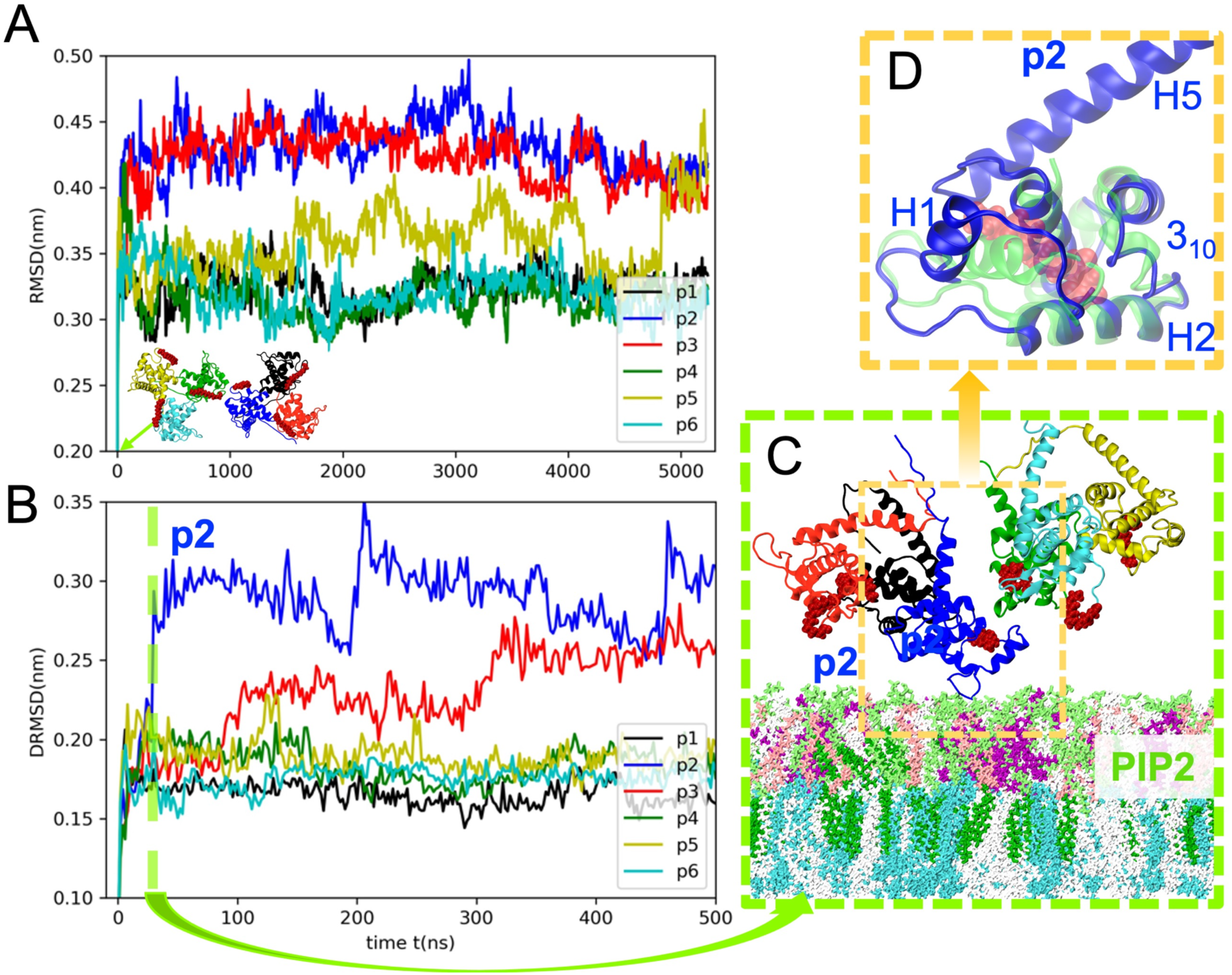
Conformational sampling of the 6 monomers in the immature MA-like dimer of trimers. (A-B) RMSD and distance RMSD (DRMSD) suggest that upon initial binding (~30ns) of p2, which initiates the binding process, the protein undergoes a structural transition that does not occur for the other monomers. (C) Side-view of the system at the instant of initial membrane anchoring of p2, (D) Structural transition of p2 (t~30ns, in blue) with respect to its initial conformation (t=0 ns; in faded green). The Myr group of p2 (t~30ns) is shown in red. Distance between H1 and 3_10_ helix increases at ~30ns; Myr group comes out of the sequestered state.

As captured by the pairwise RMSD, the p5 monomer undergoes maximum conformational fluctuation throughout the trajectory and after Myr insertion of adjacent p6 monomer (around 4600 ns), its RMSD value reaches the maximum value (0.45 nm). This correlation between Myr insertion of one monomer and conformational change of adjacent monomer requires more attention and will be analyzed, in detail, in the next section, employing time-structure based independent component analysis (tICA) method ^50–53^.

### Cooperativity of MA monomers during Myristoyl switch

Encouraged by the RMSD profile of p5 (**Figure 5A**) that captures structural transition of p5 monomer at the time of Myr insertion of adjacent p6 monomer, we aimed to explore this further by performing a dimensional reduction technique using tICA. We have considered the variable, pairwise distances between MA residues to compute the covariance matrix. A lag time of 10 ns is chosen for this purpose. MA residues in Helix 1-4 except for the N-terminal Myr group were considered in the tIC computation. The resulting tICA heatmap for p5 as 2D projections along the first and second tIC vectors (tIC1, tIC2) is shown in **Figure 6B**. The slowest relaxation modes (first two, tIC1 and tIC2) of p5 could capture the conformational switch of the MA monomer in the immature MA-like dimer undergoing a transition from unbound to the membrane-bound state. In the encounter complex, membrane binding is accomplished mainly by the HBR domain, and the Myr group is still not inserted into the membrane which occurs at the bound state. The second component (tIC2) of p5 and p6 are related to the conformational transition during the initial membrane encounter. Although neither of the first two slowest components of p5 and p6 exhibits a transition at the instant of the Myr insertion event leading to the membrane-bound state of p5 protein (~3400 ns), tIC1 of p5 shows a signature of conformational transition at the time of Myr insertion of the adjacent monomer p6 around 4700 ns (Myr dynamics data are shown in **Figure 6C**). This reveals the role of trimeric MA-MA interactions in the membrane binding process of MA protein.

**Figure 6:**
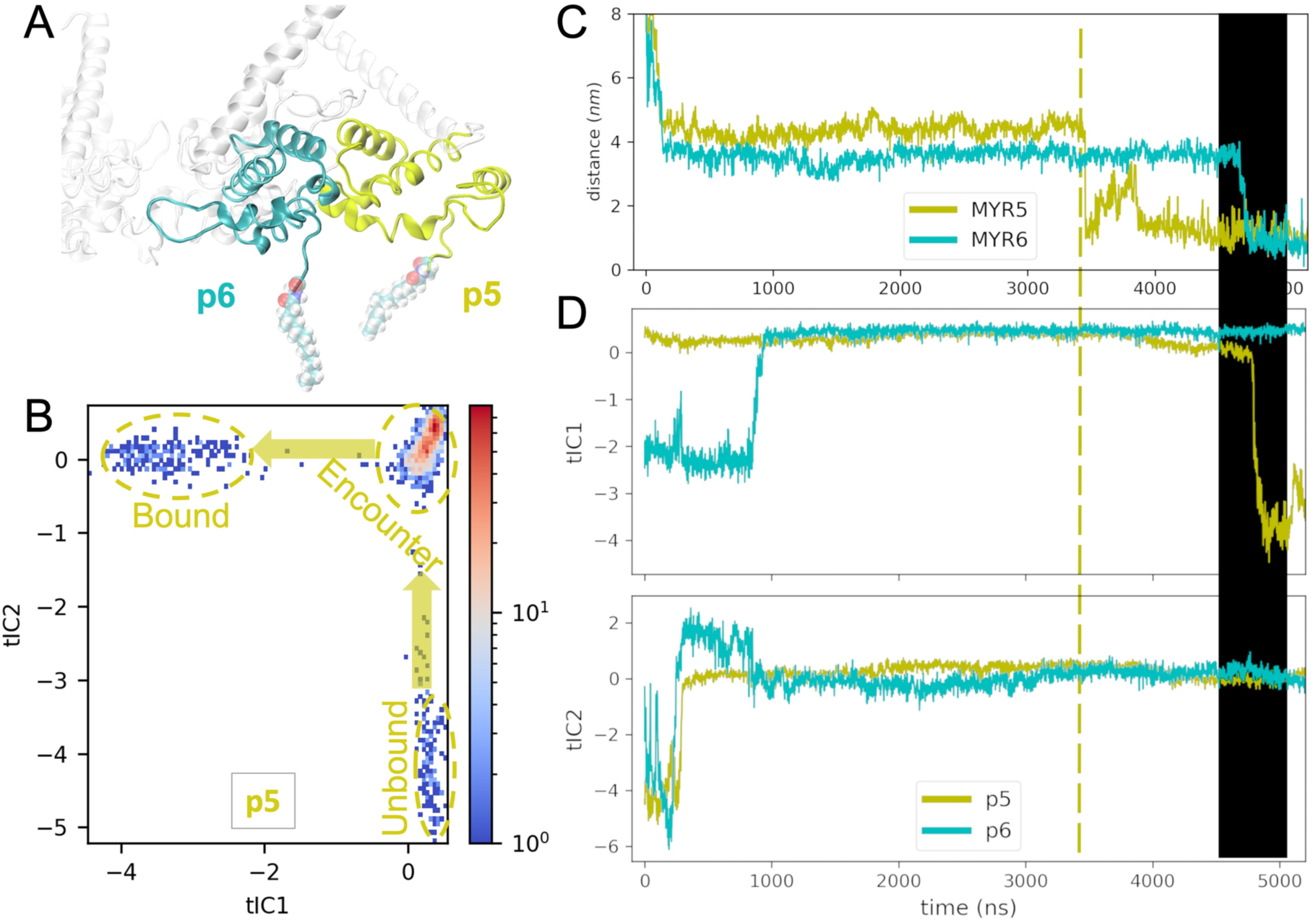
Conformational switch of MA monomer(p5) at the instant of Myr switch of an adjacent monomer(p6) is depicted with the help of two slowest time-structure based independent components (tICs) of the protein motion. (A) Snapshot of the MA trimer2 with p5,p6 at the end of the simulation, (B)2D tICA heatmap of MA monomer 5 (p5) distinguishes all the three states during membrane binding, (C) Time profile of the vertical distance of the last carbon of Myr for both p5 and p6 monomers from the bilayer center (denoted by MYR5 and MYR6 respectively). (D) Projection of the trajectory with respect to the two slowest tIC vectors. A conformational switch of p5 is observed(tIC1) at ~4700 ns during Myr insertion of p6.

### Membrane response to the initial binding of MA trimers

It has been suggested that HIV-1 Gag assembly occurs within special microdomains on the plasma membrane, the lipid composition of these domains differs notably from the rest of the host cell PM ^54–57^. These microdomains are mainly enriched with saturated lipids such as sphingomyelin (SM), cholesterol (Chol), and phosphoinositides like PI(4,5)P2 ^15, 37, 58^. This lateral heterogeneity is believed to regulate the localization and interactions of proteins and functions as cellular signaling platforms. During the HIV-1 assembly process, sphingomyelin and cholesterol in the outer bilayer form a densely packed liquid-ordered phase, called here “raft”, while PIP2 in the inner bilayer prefers a liquid-disordered phase due to its unsaturated tail. Furthermore, several studies proposed and showed that outer leaflet rafts could be trapped by inner leaflet PI/PS-Gag nanodomains through trans-bilayer coupling ^59–61^.

Here we investigate the membrane reorganization followed by initial membrane targeting of immature MA-like dimer of MA trimers. The time evolution of local lipid count data for different lipid molecules in the inner and outer leaflet are shown in **Figure 7**. After the initial encounter of all the monomers in the protein complex with the membrane (t ~ 200 ns, as marked in the dotted gray line), there is a visible increase in the number of PS and PE lipids in the inner leaflet and cholesterol, SM and PC lipids in the outer leaflet. These results help to understand Gag binding mediated remodeling of the local plasma membrane microenvironment. It has been well-accepted that the specific interaction of the Gag matrix domain and PI(4,5)P_2_ lipid plays a crucial role in the membrane binding process of matrix protein. Our simulation data suggest that MA binds to the PIP2-rich domain of the inner leaflet as the specific interactions trigger the membrane targeting process. Later, non-specific interactions with charged-lipids PS help stabilize the protein-membrane complex as seen from the increase in PS lipid count data with time (**Figure 7**). The increase in the number density of cholesterol and SM lipids on the outer leaflet is consistent with the hypothesis that the membrane binding process of matrix protein facilitates the formation of lipid rafts ^54, 62, 63^.

**Figure 7:**
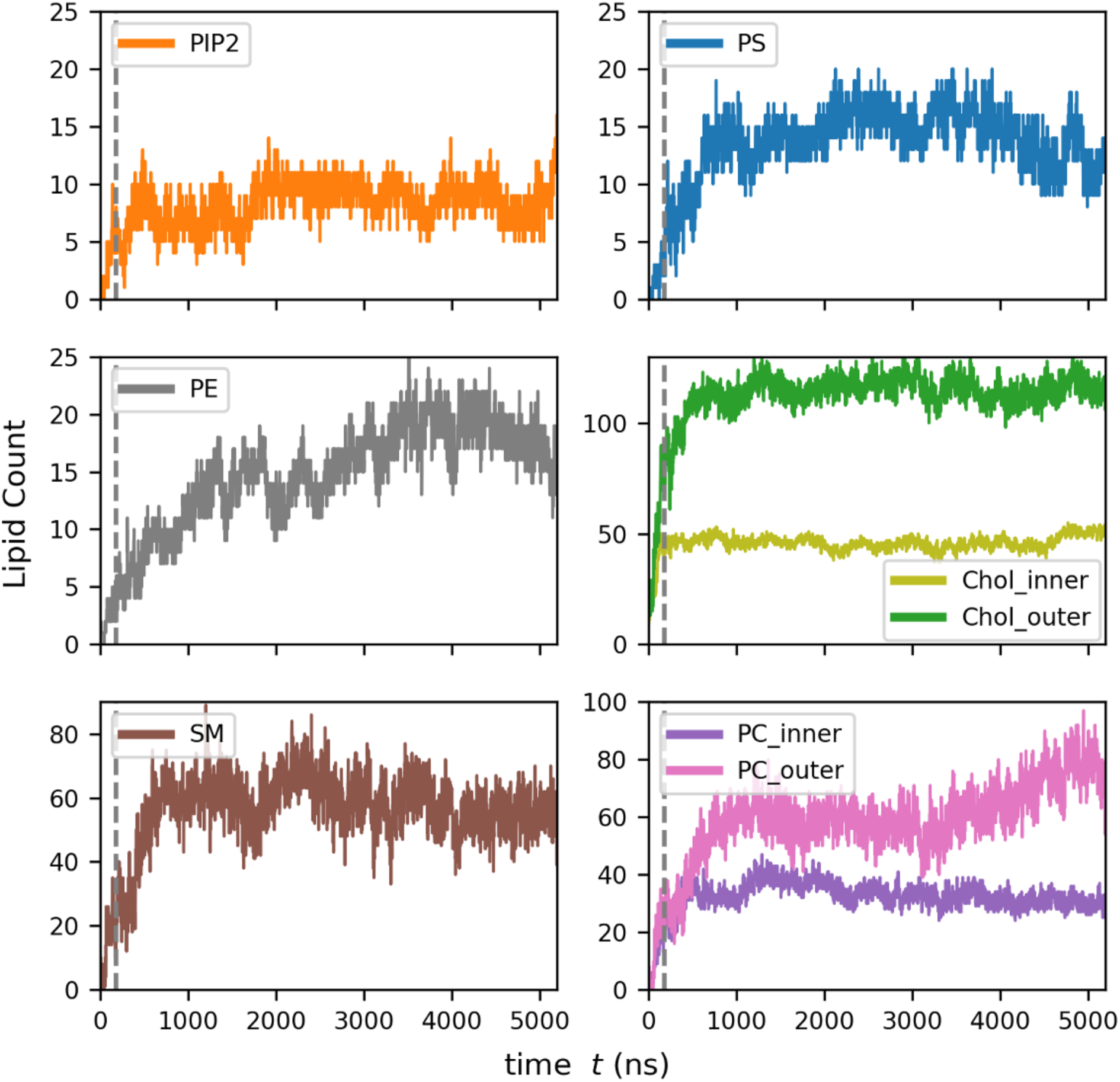
Local lipid count in the inner and outer leaflets at MA complex binding sites for the Immature MA-like dimer. Lipid density of PS, PE in the inner leaflet and Chol, SM, PC in the outer leaflet get enhanced locally. The time when all the six monomers in the complex bind the membrane is pointed by the dotted gray lines. Lipid molecule structures are shown for reference.

## Discussion and Conclusions

The present study interrogates the role of protein-protein interactions (PPI) on the Myr dynamics of the membrane-bound higher-order oligomeric structure of MA protein (a dimer of trimers). The contribution of protein-protein interactions (PPI) versus protein-lipid interactions triggering Myr exposure had yet to be understood. We have investigated the Myr insertion of the MA complex for two different systems with or without a stable trimer-trimer interface at the initial structure. Our findings reveal that the dynamics of Myr groups are impacted by the trimer-trimer interface structure. When N-terminal residues are involved in trimer-trimer interactions (TTI) in the immatureMA-like dimer, Myr groups at the trimeric interface although exposed do not show Myr insertion within simulation time (5 μs). In the random dimer model, a greater number of Myr group inserts into the membrane within the same simulation time (**Figure 2**). TTIs in these two systems are reported in **Figure 3**. This suggests that the loss of flexibility in the NTD due to the trimer-trimer interactions (like immature virion) hampers the Myr insertion process. Therefore, our simulated data support a mechanism of the Gag assembly process where membrane-bound MA forms trimers, the Myr switch happens and, finally, trimeric MA with membrane-inserted Myr groups assemble further to form dimer-of-trimers and hexamer-of-trimers.

We further analyzed the structural heterogeneity of MA monomers in the membrane-bound immature MA-like dimer and the resulting MA-lipid interactions. Unlike other monomers, the monomer that anchors the membrane first exhibits a conformational switch during membrane targeting, which was detected by DRMSD analysis (**Figure 5**). At the final step of the simulations, depending on the Myr group position, i.e., either inserted or sequestered or at the membrane surface, we observed a diverse nature of MA configurations and MA-lipid interactions (**Figure 4**). Furthermore, RMSD analysis shows interesting structural dynamics of the p5 monomer, undergoing the Myr switch. To explore this further, we have analyzed the tICA projections. The dimensionally-reduced data of MA conformations revealed the cooperativity of events (**Figure 6**). Myr insertion of one monomer is observed to induce a conformational change of an adjacent MA monomer. This correlated conformational change through the trimeric interface occurs in other trajectories as well. In this scenario, two monomers from the same trimer affect the behavior of one another. This result adds to the previous experimental prediction of Myr exposure being facilitated by the MA trimerization ^10^ and suggests the role of MA trimerization in promoting the Myr insertion event, i.e., the transition from the Myr exposed state to the membrane-inserted state. However, we do not see this conformational correlation across trimers in the 5 μs of simulation per replica.

Finally, we observe that the local lipid density has a distinctive pattern underneath the protein binding sites (**Figure 7**). In the inner leaflet, the charged PS lipid component gets enriched and stabilizes the membrane-bound encounter complex. As lipids in the inner leaflet rearrange due to the presence of the protein, lipids in the outer leaflet do it as well. Membrane binding induces an enhancement of SM and cholesterol in the outer leaflet in the regions that correspond to the protein binding sites in the inner leaflet. It should be noted that the enhancement of lipid density is measured with respect to the local lipid density MA complex encountered during membrane targeting. The increase in raft-forming lipids in the outer leaflet is reproduced in all our trajectories, both in the immature lattice as well as in the random trimer conformation.

As mentioned earlier, several factors contribute to the membrane binding of HIV-1 MA protein to the inner leaflet of the PM: (i) non-specific electrostatic interactions between the HBR of MA and anionic membrane lipids (PS, PI), (ii) specific interactions between MA protein domains and phosphatidylinositol-4,5-bisphosphate [PI(4,5)P_2_], and (iii) hydrophobic interactions between the lipidated N-terminus of MA (Myr group) and the bilayer. Although several experimental studies claimed that the myristoylation of Gag is essential for the Gag assembly process, the exact role of the Myr group is still debated. Recently, AAMD simulations by Monje-Galvan and Voth proposed that the Myr group is important for lipid sorting and membrane reorganization dynamics at the protein assembly site, yet not for PM targeting or MA monomer binding events themselves ^31^. On the other hand, NMR studies proposed that MA trimerization aids in the transition from the Myr-sequestered state to the Myr-exposed state in the solution ^10^. Later, it was shown that PI(4,5)P_2_ binding to MA induces a conformational change to facilitate Myr exposure ^18^. Our present study suggests a correlation between Myr insertion into the membrane and the conformational switch of an adjacent monomer in a MA trimer.

There is still much to be learned about MA-MA and MA-membrane interactions, lipid redistribution, membrane remodeling, and MA lattice formation during viral assembly. In our simulations, we did not obtain a membrane-bound MA complex with all the Myr residues inserted into the membrane within the simulation timescale; in fact, there was at least one Myr group remaining in a sequestered position. Further studies are therefore required to explore the dynamics of Myr insertion in MA multimeric complexes, and the factors that facilitate or hinder binding cooperativity of additional protein units as the viral assembly sites evolve.

### Simulation Model and Methods

#### Membrane model and protein-membrane system setup

Crystallographic structure of trimeric (-)Myr-MA was retrieved from PDBID:1HIW in the protein data bank (https://www.rcsb.org) ^7^. The monomeric (+)Myr-MA structure (PDB ID: 1UPH) ^10^ with all MA residues starting from residue 1 (GLYM) to residue 131(TYR) was then superimposed to the trimer 1HIW to obtain the final trimeric (+)Myr-MA structure in the immature lattice conformation ^12^. For the random dimer simulations, the PDB structure of trimeric MA (PDB ID:1HIW) was lipidated using the CHARMM_GUI webserver and used. Details of PPI in these systems are provided in **Figure 1** and **Figure 3**.

The membrane model designed for the study was based on lipidomics analysis of HIV-1 particles reported by Lorizate *et al*. ^36^ **Table 1** contains the molecular fraction of each lipid component per leaflet. This model mimics the asymmetric nature of the PM and allows us to examine the effect of trans-bilayer interactions that results upon protein binding. The asymmetric bilayers were initially built as two symmetric bilayers, each with the corresponding lipid composition for the desired asymmetric bilayer. The symmetric membrane models were equilibrated for 100ns before merging one leaflet per model. Given the difference in lipid content, each leaflet had a different surface area. The smaller leaflet patch, modeling the outer leaflet of the PM, was replicated to match the surface area of the model for the inner leaflet. Molecules outside of the larger leaflet patch, modeling the inner leaflet of the bilayer, were deleted to render the asymmetric model. This system was further equilibrated for 250ns before merging the equilibrated protein coordinates at 1.0 nm above the bilayer using VMD software ^64^. Fully hydrated bilayers and an MA trimer were built and equilibrated separately using CHARMM-GUI Membrane Builder and Quick-Solvator, respectively ^65–69^. For immature MA-like dimer simulations, initially, two MA trimers are arranged where N-terminal residues, H1 and 3_10_ helices of a MA monomer of trimer1 are in close proximity to the same protein domains of another MA monomer of trimer2 (as reported by recent cryoET experiment for MA lattice in immature virions ^12^. All systems were rendered electrostatically neutral using a 0.15 M KCl salt solution. The total size of the immature MA-like dimer systems was ~ 590,000 atoms, with a box size of 16.3 nm × 16.5 nm × 22.3 nm. The random dimer system size was ~ 490,000 atoms and the box size was 18.6 nm. Two replicas were run for 5 μs for each system, for a total of 20 μs.

**Table 1:** Lipid content for the membrane model used in this study.

|  | **Lipid species** | DOPC | LSM | Chol | PIP2 | DOPS | POPE |
| --- | --- | --- | --- | --- | --- | --- | --- |
| **Lipid fraction per leaflet** | **Inner Leaflet** | 0.4 | 0.0 | 0.15 | 0.15 | 0.15 | 0.15 |
|  | **Outer Leaflet** | 0.3 | 0.35 | 0.35 | 0.0 | 0.0 | 0.0 |

#### AAMD simulation settings and trajectory analysis

All simulations used the CHARMM36m force field ^70^; equilibration runs were performed in GROMACS 2019 ^71^, and micro-second production runs were carried out on the ANTON2 machine ^72^. Energy minimization was performed using the steepest descent integrator until the force was reduced to 1000 kJ mol^-1^ nm^-1^. The initial relaxation of each system was carried out using the default steps obtained from CHARMM-GUI, the platform used to build the initial simulation coordinates. The default relaxation scripts run first in the constant NVT ensemble and then in the constant NPT ensemble for a total of 2 ns. During this phase, the Cα backbone of the protein is harmonically restrained with a force constant of 1000 kJ mol^-1^ nm^-2^. Afterward, constant NPT dynamics were used for the equilibration and production runs without any positional restraints on the system. The temperature was kept constant at 310.15 K using the Nose-Hoover thermostat with a 1.0 ps coupling constant ^73, 74^. The pressure was set at 1 bar and controlled using the Parinello-Rahman barostat semi-isotropically due to the presence of membrane, the compressibility factor was set at 4.5 ×10^-5^ bar^-1^ with a coupling time constant of 5.0 ps ^75, 76^. Throughout the trajectories in GROMACS, van der Waals interactions were computed using a force-switching function between 1.0 and 1.2 nm, long-range electrostatics were evaluated using Particle Mesh Ewald ^77^, and hydrogen bonds constrained using the LINCS algorithm ^78^.

All systems were run for at least 200ns in GROMACS prior to transferring them to the Anton2 machine. Simulation parameters for the Anton2 runs were set by internal ark guesser files; as such, the cut-off values to compute interactions between neighboring atoms are selected automatically during system preparation. Long-range electrostatics in this machine were computed using the Gaussian Split Ewald algorithm ^79^, and hydrogen bonds constrained using the SHAKE algorithm ^80^. Finally, the Nose-Hoover thermostat and MTK barostat controls are used to run NPT dynamics using the Multigrator integrator algorithm on Anton2 ^81^.

Analysis of the trajectories was performed using GROMACS, MDAnalysis ^82, 83^, MDTraj^84^ and VMD Tcl scripts^64^. Time-lagged independent component analysis (tICA) ^50–53^ was run as a dimensionality reduction technique to quantify the protein conformational change during membrane binding; this analysis was done using MDTraj and MSMBuilder ^85^.

## Acknowledgments

This research was supported by the National Institute of Allergy and Infectious Diseases (NIAID) for the Behavior of HIV in Viral Environments (B-HIVE) Center of the National Institutes of Health (NIH grant U54 AI170855) and the National Institute of General Medical Sciences (NIH grant R01 GM063796). Computational resources were provided by the Pittsburgh Super Computing Center through the Anton2 machine. Part of this work was also completed with resources from the University of Chicago Research Computing Center, the Extreme Science and Engineering Discovery Environment (XSEDE), and the NIH-funded Beagle-3 computer (NIH award 1S10OD028655-01).

